# Biochemical and Biophysical Characterization of the dsDNA packaging motor from the *Lactococcus lactis* bacteriophage asccphi28

**DOI:** 10.1101/2020.11.17.383745

**Authors:** Emilio Reyes-Aldrete, Erik A. Dill, Cecile Bussetta, Michal R. Szymanski, Geoffrey Diemer, Priyank Maindola, Mark A. White, Wlodzimierz M. Bujalowski, Kyung H. Choi, Marc C. Morais

## Abstract

Double-stranded DNA viruses package their genomes into pre-assembled protein procapsids. This process is driven by macromolecular motors that transiently assemble at a unique vertex of the procapsid and utilize homomeric ring ATPases to couple genome encapsidation to ATP hydrolysis. Here we describe biochemical and biophysical characterization of the packaging ATPase from Lactococcus lactis phage asccφ28. Size-exclusion chromatography, analytical ultracentrifugation, small angle x-ray scattering, and negative stain TEM indicate that the ~45 kDa protein formed a 443 kDa cylindrical assembly with a maximum dimension of ~155 Å and radius of gyration of ~54 Å. Together with the dimensions of the crystallographic asymmetric unit from preliminary X-ray diffraction experiments, these results indicate that gp11 forms a decameric D5-symmetric complex consisting of two pentameric rings related by 2-fold symmetry. Additional kinetic analysis shows that recombinantly expressed gp11 has ATPase activity comparable to that of functional ATPase rings assembled on procapsids in other genome packaging systems. Hence, gp11 forms rings in solution that likely reflect the fully assembled ATPases in active virus-bound motor complexes. Whereas ATPase functionality in other dsDNA phage packaging systems requires assembly on viral capsids, the ability to form functional rings in solution imparts gp11 with significant advantages for high resolution structural studies and rigorous biophysical/biochemical analysis.

## 1. Introduction

Tailed bacteriophages have historically served as model systems for mechanistic investigations of powerful molecular motors that use ATP hydrolysis to translocate double-stranded DNA into pre-formed virus protein capsids [1–4]. Most large ssDNA viruses and dsDNA viruses—such as herpes virus, pox virus, adenovirus, and all the tailed dsDNA bacteriophages—assemble an empty virus shell, and then actively translocate DNA into this procapsid. There are considerable entropic and enthalpic costs associated with compacting DNA to near-crystalline densities inside virus shells [3,5,6]. DNA must be bent significantly and repeatedly, and it must be condensed without any knots, tangles, or other topological anomalies that would prevent genome ejection during subsequent infection events. To package DNA efficiently, viruses code for homomeric ring ASCE ATPases that tightly couple ATP hydrolysis and DNA translocation [1,3,4,7,8]. This ancient ATPase superfamily is ubiquitous across all three domains of life, and its members are typically involved in polymer manipulation tasks (e.g. protein degradation, chromosome segregation, DNA recombination, DNA strand separation, and conjugation), or in molecular segregation such as proton movement by the F1-ATP synthase [9–12]. Thus, understanding the mechanistic principles underlying viral DNA packaging will also illuminate the mechano-chemistry of a broad class of molecular motors responsible for basic macromolecular partitioning processes. Within the ASCE NTPase superfamily, virally encoded dsDNA packaging motors are especially powerful, capable of producing forces in excess of 50 pN [13,14]. Hence the virally encoded ATPases that drive dsDNA encapsidation provide a unique window into how maximum mechanical force can be extracted from chemical hydrolysis of ATP. In addition to serving as model systems to investigate the biophysical challenges associated with DNA condensation and ATPase mechanochemistry, these motors are attractive targets for therapeutics that inhibit dsDNA virus proliferation since there is no direct counterpart in eukaryotic cells [15].

Given the importance of viral dsDNA packaging motors in both applied and basic biomedical research, considerable effort has been exerted to understand the molecular basis of genome packaging. As a result, several different bacteriophage dsDNA packaging systems have been developed to experimentally interrogate the encapsidation process, each of which have relative advantages and disadvantages [1–3]. Common to all systems, the packaging motors include a portal/connector protein that provides the “portal” for DNA entry and exit, and an ATPase that powers DNA translocation through the portal and into the capsid. While the packaging motors in each system essentially perform the same task, use similar molecular machinery, and likely share common underlying mechano-chemistry, there are important differences which must be considered. To provide a simple conceptual framework for categorizing packaging motors, we distinguish between two different types: 1) those that must cut their DNA at the beginning and end of packaging to encapsidate concatenated genomes; and 2) those that simply package a fixed unit-length genome.

Two of the most well-known examples of genome-cutting packaging systems are bacteriophages lambda and T4 [1,2]. In these phages, a concatenated multiple-copy genome is transcribed, and thus the motor is not simply a translocating machine. It first must find the genome start site on the concatenated multi-genome, make a cut in the genome, and only then begin packaging the DNA into the capsid. Further, as the capsid fills with DNA, the motor must somehow sense that packaging is complete and cut the genome again such that the motor-DNA complex can detach from the filled head and re-attach to another empty procapsid. In bacteriophage lambda a specific DNA sequence known as a “cos” site is recognized as the cut site at the beginning and end of packaging. Bacteriophage T4 uses a “headful” mechanism, wherein the phage somehow recognizes that slightly more than one genome-length of DNA has been packaged, thus filling the procapsid.

To accomplish these additional tasks, genome-cutting motors require additional protein components to identify the genome and locate the appropriate start site for packaging, as well as a functional nuclease to cut the genome at the beginning and the end of packaging. Hence, genome-cutting phages also code for a fully functional RNaseH-like nuclease that is fused to the ATPase as a C-terminal domain and which cuts DNA at the beginning and end of packaging. Due to this nucleolytic function, these ATPases are often referred to as large terminases. Genome-cutting systems also code for a so-called small terminase protein that is likely involved in genome recognition. Despite adopting the “terminase” nomenclature, no enzymatic functions are associated with the small terminase. Neither the exact function of the small terminase, nor how it is incorporated in the motor complex, is well understood.

Fixed length packaging systems are somewhat simpler. In these types of phages, single unitlength genomes are produced rather than the concatenated multi-genomes utilized by genome-cutting phages. Because the polymerases in these phages use a protein primer to initiate replication, the genomes of fixed length phages have a protein covalently attached to the 5’-end of each strand of genomic DNA. The ATPase assembles at the portal vertex, engages its DNA substrate by recognizing and binding the terminal protein at a genomic 5’-end, packages until the entire nucleic acid is encapsulated, and then detaches, possibly to assemble at the portal vertex of another empty capsid. The most thoroughly characterized fixed length packaging system is the bacteriophage φ29 system. Since φ29 packages a unit length genome, its packaging motor is accordingly simpler. There is no need for a nuclease function to cut the DNA, and thus the φ29 ATPase lacks a fully functional nuclease domain and is only ~ 60% the size of ATPases/large terminases in genome-cutting packaging systems [16]. Similarly, the portal protein is only ~60% of the size of portals in cutting systems [17], possibly because it does not have or need any structural components to sense that the capsid is full and/or transmit this signal to the rest of the motor. Further, the 5’-terminal protein provides a unique identifying feature of genomic DNA, and thus these phages do not use a small terminase for genome recognition. Thus, φ29 is in some sense a stripped-down version of the genome-cutting packaging motors. However, φ29 is by no means simple; structural and single molecule results indicate that the φ29 packaging motor operates in a highly coordinated manner to regulate the activity of its subunits and to efficiently generate force to translocate DNA [18–21]. Additionally, φ29 is unusual in that it requires a virally encoded structural RNA molecule (pRNA) that acts as a scaffold for assembly of a functional ATPase ring, and which is not present in headful packaging systems [17,22–25]. Thus, neither the genome-cutting nor fixed genome-length systems represent the universal essential minimum necessary for operation of a packaging motor.

An additional complicating factor in all current model systems is that in both genome-cutting and unit-length packaging motors, the ATPases form functional rings only by virtue of assembling on the capsid (as in T4) [26], on the small terminase (as in lambda) [1, 27], or on the pRNA (as in φ29) [17,25]. As a result, rigorous biophysical studies to interrogate inter-subunit interactions are challenging, and structural information regarding the functional ATPase assembly has thus far been limited to fitting X-ray structures of various motor components into low resolution cryoEM maps of motor complexes assembled on procapsids [26,28]. Understanding the molecular basis of subunit coordination would thus be facilitated by the availability of stable ATPase ring assemblies that can be interrogated biophysically and via atomic resolution structural information to learn how the ATPase subunits interact to efficiently translocate DNA.

It would thus be useful to develop a packaging system that is like φ29 in that it has fewer, smaller, and simpler motor components, but which does not need a pRNA to assemble a functional ATPase ring. Such a motor would reflect the most experimentally useful aspects of both types of systems and likely represent the bare minimum necessary to translocate dsDNA. Toward this end, we have characterized the DNA packaging ATPase from the *Lactococcus lactis* phage asccφ28 [29]. Like φ29, asccφ28 packages a unit-length genome that has a terminal protein covalently attached to each 5’-end of its genomic DNA. Hence, asccφ28 also codes for smaller, simpler versions of the portal protein and the ATPase, similar to φ29. However, there is no evidence for a pRNA in its genome sequence. Since a primary function of the pRNA in φ29 is to act as a scaffold for assembly of a functional ATPase ring [3,17,23,28], we hypothesized that the ATPase from asccφ28 might have evolved to form functional rings on its own. If so, the ATPase from φ28 would be an excellent model for rigorous biophysical and high-resolution structural analysis of functional ATPase ring motors that drive viral dsDNA genome packaging.

Indeed, using a combination of size-exclusion chromatography (SEC), analytical ultra-centrifugation (AUC), small X-ray scattering (SAXS), negative-stain transmission electron microscopy (TEM), and preliminary X-ray crystallographic diffraction, we show that recombinantly expressed gene product 11 (gp11) from asccφ28 is a highly soluble and stable decameric assembly consisting of two pentameric rings related by D5 point group symmetry. Additional kinetic characterization of NTPase binding and hydrolysis shows that the assembly binds and hydrolyzes ATP similarly to the ATPases in φ29 and other phages, but only once the ATPases in these other systems have assembled functional rings on the procapsid. These results suggest that the pentameric rings in the D5 decamer reflect the biological assembly of the gp11 packaging ATPase on asccφ28 procapsids. Hence, the dsDNA packaging motor gp11 from bacteriophage asccφ28 provides a unique opportunity to examine a phage dsDNA-packaging motor apart from other packaging components. Studying an isolated, functional ATPase ring motor will facilitate rigorous biophysical analysis, more straightforward kinetic analysis, and high-resolution structure determination, expediting a comprehensive understanding of viral dsDNA packaging motors and the mechano-chemistry common to the larger superfamily of ASCE ATPases to which they belong.

## 2. Materials and Methods

### 2.1 Cloning

A BLAST search identified phage asccφ28 gene product 11 (gp11) as orthologous to the bacteriophage φ29 packaging ATPase, with 45% sequence similarity and 28% amino acid identity [30]. A recombinant, codon-optimized gene for gp11 was synthesized by DNA 2.0 (now ATUM) based on the published sequence of phage asccφ28 (GenBank ascension number EU438902) [29]. The synthesized gene was inserted into a pET-30a(+) vector (Novagen) with kanamycin resistance using NdeI and XhoI restriction sites, resulting in a final construct with an additional C-terminal, - LEHHHHHH tag to facilitate purification by metal-affinity chromatography [31]. DNA sequencing confirmed that the final construct matched the published amino acid sequence of asccφ28 gp11.

### 2.2 Protein expression and purification

*E. coli* cells transformed with the synthetic construct were grown in Luria-Bertani medium with 30 μg/ml of kanamycin at 37°C until optical density at 600 nm reached ~0.6 (1 cm path). The BL21 (DE3) strain of *E. coli* (Novagen) was predominantly used, but other strains also show good expression. Maximum yield of soluble protein was obtained by induction with isopropyl-β-D-1-thiogalactopyranoside (IPTG) at a final concentration of 0.1 mM and expression overnight at 18°C. Cells were harvested by centrifugation at 4000 g for 20 min at 4°C. Cell pellets were then either stored at −20°C or immediately resuspended in a lysis/extraction buffer containing 50 mM Na2HPO4, 1M NaCl, 5 mM β-mercaptoethanol, and 5 mM imidazole. The typical volume of lysis/extraction buffer used was 25 mL for a pellet harvested from 1 L of cell culture. Resuspended cells were kept on ice and lysed by sonication with the Microson XL200 Ultrasonic Cell Disruptor (Misonix). The sonication regime involved 30 s pulse treatments with 30 s pauses between pulses until the mixture appeared completely homogenous (typically after ~15-20 pulse treatments). The soluble protein fraction was then separated from cell lysate solids by centrifugation at 17,000 g for 30 min at 4°C. Immobilized metal affinity chromatography was used for the first purification step. The supernatant containing the soluble, 6xHis-tagged protein fraction was incubated for thirty minutes at 4°C with Talon resin (Clontech Laboratories/Takara Bio USA) that had been previously equilibrated in lysis/extraction buffer, typically employing ~1 mL of resin for ~25 mL of supernatant. The resin-bound protein was then transferred to a chromatography column and eluted in 1 mL fractions with a linear gradient going from 5 mM to 150 mM imidazole. Talon column fractions that contained gp11 were then pooled, concentrated by ultrafiltration, and further purified by size exclusion chromatography on a HiLoad Superdex 200 gel filtration column (GE Healthcare). The column was equilibrated and run with a buffer containing 50 mM sodium phosphate, pH 8.1, 400 mM sodium chloride, and 1 mM dithiothreitol. Fractions from the single elution peak were assessed by SDS-PAGE and combined for subsequent characterization.

### 2.3 Size estimation by gel filtration chromatography

Gel filtration chromatography was used to estimate the protein molecular weight by comparing gp11’s elution profile with protein standards of known molecular weight. A commercial standard containing 5 different molecular weight markers (Biorad) was run on the same HiLoad Superdex 200 16/60 column, in buffer containing 400 mM NaCl, 25 mM Tris pH 8.0, and 1 mM DTT. The standards in the mixture were Thyroglobulin (670 kDa), γ-globulin (158 kDa), ovalbumin (44 kDa), myoglobin (17 kDa), and Vitamin B12 (1.35 kDa). The elution volume for the peak of each standard was used to plot a standard curve, and the elution volume of gp11 was plotted on this curve to estimate the molecular weight.

### 2.4 Analytical ultracentrifugation

Purified protein was dialyzed into a buffer containing 20 mM Tris-HCl, pH 8.1 at 20°C, 400 mM NaCl and 5 mM β-ME and subsequently centrifuged at 27000 g for 20 min at 4°C. Concentration was determined spectrophotometrically based on Edelhoch’s method, with a predicted extinction coefficient ε280 = 42400 cm^-1^M^-1^ for the 44.6 kDa monomer [32,33]. Analytical ultracentrifugation experiments were performed with an Optima XL-A analytical ultracentrifuge (Beckman Inc., Palo Alto, CA), equipped with absorbance optics and An60Ti rotor; all experiments were carried out at 20° C. Absorbance data were collected by scanning the sample cells at wavelength intervals of 0.001 cm in the step mode with 5 averages per step. Each experiment was conducted at two or three rotor speeds, starting with the lowest and finishing with the highest rotor speed. The sedimentation was assumed to be at equilibrium when consecutive scans, separated by intervals of 8 hours, did not indicate any change.

### 2.5 Solution small-angle X-ray scattering

SAXS experiments were performed at the ALS beamline 12.3.1 (SIBYLS; Lawrence Berkeley National Laboratory Berkeley, CA, USA). Scattering intensities I(q) for protein and buffer samples were recorded as a function of scattering vector q (q = 4πsinθ/Λ, where 2θ is the scattering angle and Λ is the X-ray wavelength). The sample-to-detector distance was set to 1.5 m, which resulted in a q range of 0.01 - 0.32Å^-1^, Λ was 1.0 Å and all experiments were performed at 20 °C. The data collection strategy described by Hura was used in this study [34]. Briefly, SAXS data were collected for three protein concentrations (1.4, 2.0, and 2.7 mg/ml) and for two buffer samples. For each sample measurement, SAXS data were collected for three X-ray exposures – one long exposure (10 s) flanked by two short exposures (1 s) to assess radiation damage. Buffer scattering contributions were subtracted from sample scattering data using the program ogreNew (SIBYLS beamline). Data analysis was performed using the program package PRIMUS from the ATSAS suite 2.3 [35,36]. Experimental SAXS data obtained for different protein concentrations were analyzed for aggregation and folding state using Guinier and Kratky plots, respectively. The forward scattering intensity I(0) and the radius of gyration RG were evaluated using the Guinier approximation: I(q) ≈ I(0) exp(−q2R_g_)^2^ / 3, with the limits qR_g_ < 1.3. These parameters were also determined from the pair-distance distribution function P(R), which was calculated from the entire scattering patterns via indirect Fourier inversion of the scattering intensity I(q) using the program GNOM [37]. The maximum particle diameter D_max_ was also estimated from the P(R). The hydrated volume VP of the particle was computed using the Porod equation: V_P_ = 2*π*^2^I(0)/Q, where I(0) is the extrapolated scattering intensity at zero angle and Q is the Porod invariant [38,39]. The molecular mass of a globular protein can then be estimated from the value of its hydrated volume [39]. The overall shape of the protein was modeled *ab initio* by fitting the SAXS data to the calculated SAXS profile of a chain-like ensemble of dummy residues in reciprocal space using the program GASBOR, version 2.2i [40]. Ten independent calculations were performed with a D5 symmetry restriction (see below for choice of symmetry).

### 2.6 Electron microscopy

A continuous carbon-film copper grid (Electron Microscopy Sciences) was plasma cleaned for 30 s. 3 μL of purified protein at 0.5 mg/ml was applied to the grid for 3 s then blotted. 3 μL of 1% uranyl acetate (aq) stain was applied three times and blotted after each application. Grids were imaged on a JEM 2100 microscope equipped with a Gatan US4000 CCD camera.

### 2.7 Crystallization

Protein in buffer containing 50 mM sodium phosphate, pH 8.1, 400 mM sodium chloride, and 1 mM dithiothreitol was concentrated via filtration to ~4.5 mg/ml. Several commercial sparse matrix screens, including the Wizard Classic screen (Rigaku) and Salt RX screen (Hampton Research), were used to determine initial crystallization conditions using an automated PHENIX crystallization robot and Intelliplate 96-well sitting drop trays (both from Art Robbins Instruments). Promising crystals were screened on a Rigaku FR-E++ X-ray generator with an RAXIS-IV++ crystallography system. Subsequent optimization screens were set up manually at various temperatures in 24 well VDX trays using hanging-drop vapor-diffusion geometry.

### 2.8 X-ray diffraction data collection and processing

Seleno-methionine derivatized gp11 crystals grown from two distinct optimized conditions (A and B) were shipped to Argonne National Laboratory, and data collected at the Advanced Photon Source beamline 21 ID-F on a MAR 225 CCD detector. Prior to freezing in liquid nitrogen, samples of crystal form A were soaked in well solution supplemented with 20% glycerol to prevent formation of crystalline ice. One microliter of cryoprotectant solution was added directly to the crystallization drop and allowed to soak for 10-30 minutes prior to freezing. Data were collected with an oscillation angle of 1.0° and one image per second, then indexed, integrated, merged and scaled with HKL-3000 [41]. Crystal form B was directly frozen in liquid nitrogen; citrate in the crystallization buffer was of sufficient concentration to act as a cryoprotectant. Data were collected using the helical scan technique, translating the crystal through the x-ray beam to reduce radiation damage. Images were collected every 0.5 s in 0.5° steps.

### 2.9 Activity assays

NTPase activities for gp11 were determined using a continuous coupled assay with the enzymes pyruvate kinase and lactate dehydrogenase. The reaction scheme (figure 7a) involves uptake by pyruvate kinase of the NDP generated by the NTPase, and transfer of the phosphate group from phosphoenolpyruvate (PEP) to regenerate NTP and pyruvate. The released pyruvate is then taken by the enzyme lactate dehydrogenase, along with reduced nicotinamide adenine dinucleotide (NADH), to generate lactate and oxidized nicotinamide adenine dinucleotide (NAD^+^). The progression of the reaction is monitored by measuring the reduction in absorbance at 340nm as NADH is converted to NAD^+^. The reaction mixture contains 10 U/mL rabbit muscle pyruvate kinase, 15 U/mL rabbit muscle lactate dehydrogenase, 2mM PEP, and 0.3mM NADH. Reactions were carried out in a 50mM Tris, 72mM KCl, 7.2mM MgSO_4_ (TKM) buffer as first described in [42]; all reagents were purchased from Sigma Aldrich. Data was analyzed as the simplest case described by the Michaelis-Menten equation where NTP is rate limiting, and the steady-state rate of ATP hydrolysis, v, is given by equation 1.

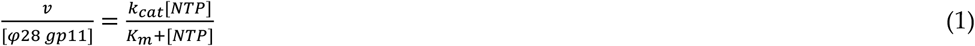

## 3. Results

### 3.1. Identification of the asccφ28 ATPase

Bacteriophage φ29 has long served as a model system for investigation of genome packaging in dsDNA viruses. However, attempts to crystallize its full-length dsDNA packaging ATPase have not been successful. A BLAST search conducted for orthologs of the φ29 DNA packaging ATPase returned only eight closely related constructs (25-77% amino acid identity) that both belonged to dsDNA phages and were predicted to have an ATP-binding fold [30]. To identify potential crystallization targets, these eight sequences were run through the XtalPred server [43]. The server returned a classification of “optimal” for only one of the eight pre-selected BLAST hits, belonging to the putative DNA packaging ATPase from the *Lactococcus lactis* phage asccφ28 (φ28), which has a 55.6% Smith-Waterman sequence similarity score to the φ29 ATPase [44]. Phage asccφ28 was identified as an infectious agent of dairy fermentation strains of *Lactococcus lactis* [29]. Morphological characterization and genomic analysis carried out in this same study indicated that the newly discovered phage asccφ28 is genetically comparable to bacteriophage φ29. Gross similarities to φ29 include genome size (19-20 kbp), prolate icosahedral capsid size and morphology, and a short non-contractile tail. Additional important similarities include packaging a unit-length genome with a terminal protein covalently attached to each 5’-end and smaller versions of the portal and ATPase motor components, as compared with other phages.

### 3.2 Cloning, expression and purification

A recombinant form of open reading frame 11, the putative ATPase from asccφ28, was expressed in E. coli BL21(DE3). A large amount of soluble protein was obtained after induction with 1 mM IPTG and expression at 18°C overnight. A first purification step with Talon resin in a 50 mM sodium phosphate and 300 mM sodium chloride buffer yielded late-eluting fractions which were over 95% pure as judged by SDS-PAGE (Figure 1) [45]. A second purification step was carried out by size-exclusion chromatography in an AKTA FPLC system equipped with a High Load 16/60 Superdex 200 column, resulting in greater than 95% purity. LCMS-mass spec analysis of tryposin-proteolyzed gel bands confirmed the purified protein was gp11.

**Figure 1:**
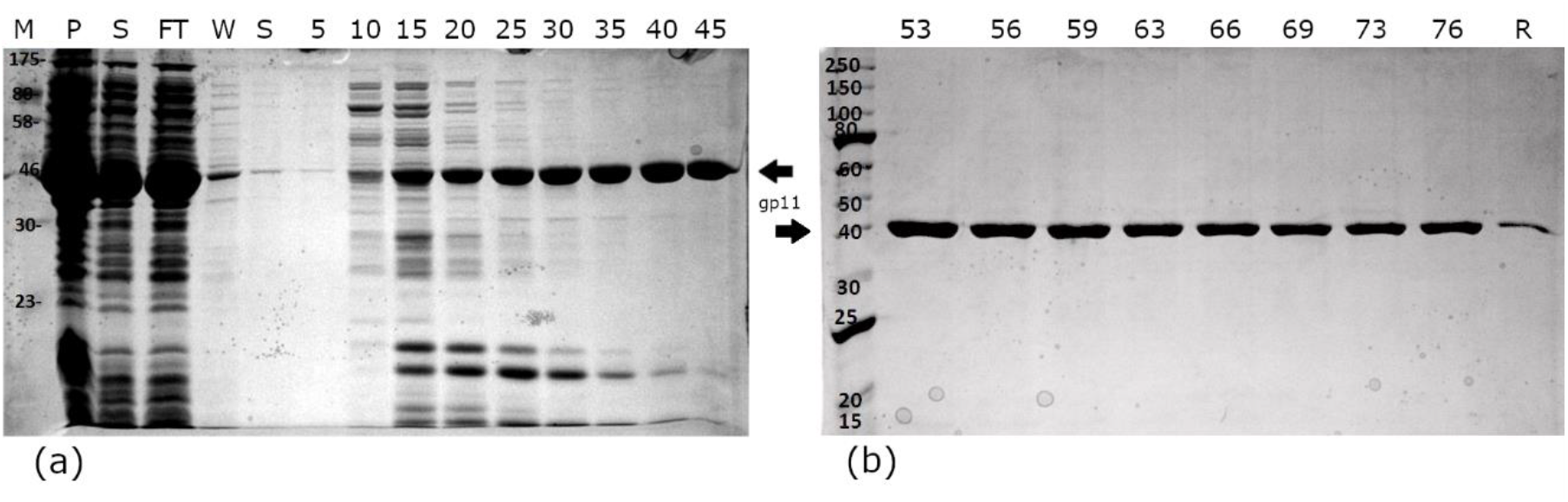
Metal-affinity purification of gp11 protein from cell lysate. SDS-PAGE gels show the early (a) and late (b) eluting fractions. Although the late-eluting fractions appeared to contain a single species, size-exclusion chromatography was used as a second purification step to increase purity. Column labels M – Molecular weight standard, P – Pellet (insoluble material from cell lysis), S – Soluble material from cell lysis, FT – Flow through from Talon resin binding, W – Wash of resin. Numbering references fractions eluted from the column. R – Residual protein remaining bound to the Talon resin

### 3.3 Size estimation by gel filtration chromatography

The chromatographic profile obtained during gel filtration suggested that the protein exists as a large assembly. Gp11 has a calculated molecular mass of 44.6 kDa but eluted from the Superdex 200 gel filtration column near the beginning of the elution profile, suggesting a larger molecular weight. Comparison to a typical elution profile of standard proteins in the same chromatographic medium and flow rate pointed towards an apparent molecular weight between that of IgG (158 KDa) and Ferritin (440 KDa) (figure 2), suggesting that gp11 had assembled a higher order oligomer with between ~4 and ~10 copies. Similarly, while SDS-PAGE bands were consistent with a 45 kDa protein, native PAGE again showed a higher apparent molecular weight (results not shown).

**Figure 2:**
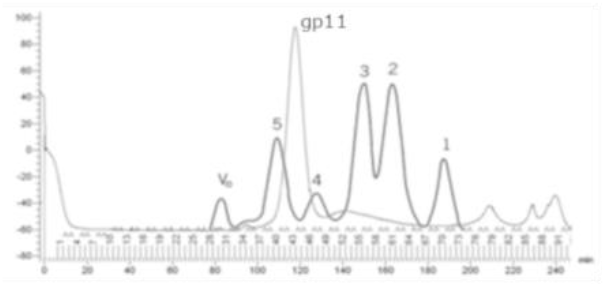
Light gray line is a typical elution profile from a Superdex 200 (16/60) gel filtration column for gp11-containing fractions combined after metal-affinity purification. Dark gray overlay is the elution profile of a standard protein mixture over the same flow rate conditions. Standards are 1. Myoglobin, Mr 17,000 Da; 2. Ovalbumin, Mr 43,000 Da; 3. Albumin, Mr 67,000 Da; 4. IgG, Mr 158,000; 5. Ferritin, Mr 440,000.

### 3.4 Analytical ultracentrifugation

To determine the molecular weight of the assembly observed in gel filtration and native gel electrophoresis, we carried out analytical ultracentrifugation measurements. Sedimentation equilibrium experiments were performed for multiple rotor speeds (figure 3). The smooth curves overlying the data are simulations using the best fit parameters resulting from a global NLLS with Mi and b as fitting parameters. For the n-component system, the total concentration at radial position r, c_r_, is defined by:

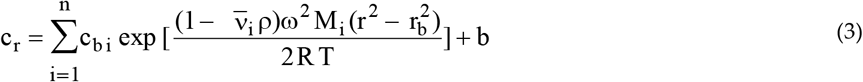

where c_bi_, v_i_, and M_i_ are the concentration at the bottom of the cell, partial specific volume, and molecular mass of the “I” component, respectively; ρ is the density of the solution, ω is the angular velocity, and b is the base-line error term. Partial specific volume of gp11 (v_i_) was calculated from the amino acid composition of the protein according to Lee and Timasheff [46] and was 0.738 mL/g. The molecular mass calculated from the amino acid composition of gp11 is 44,585 Da. Molecular mass for the sample was 443,472 ± 8000 Da which is consistent with a decameric assembly of gp11. Sedimentation velocity experiments also indicated a single species corresponding to a decamer (data not shown).

**Figure 3:**
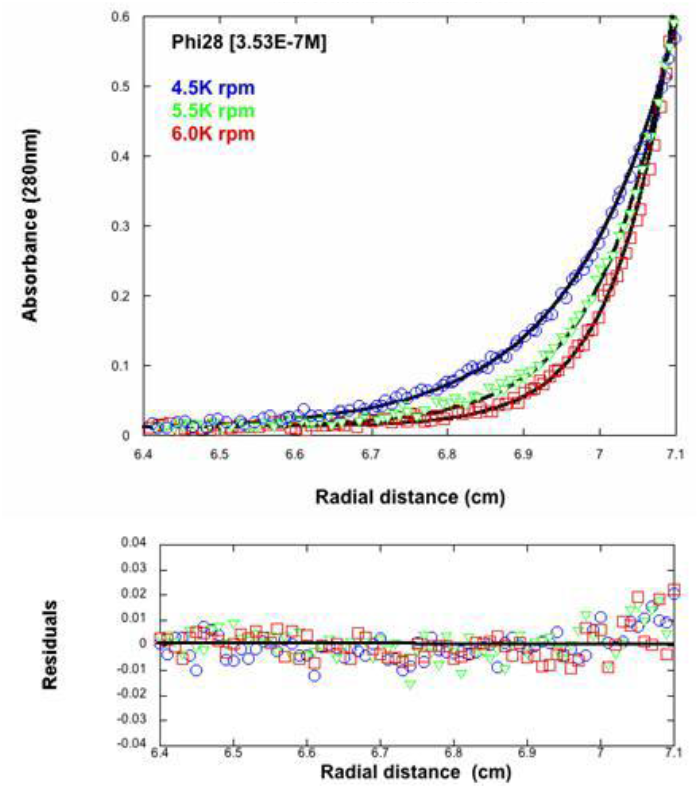
Sedimentation equilibrium centrifugation of gp11. Rotor speeds were 4.5, 5.5, and 6.0 kRPM. Protein concentration was 0.353 μM (0.016 mg/mL). In solution, gp11 has an apparent molecular weight of 443 kDa, about ten times the monomeric weight.

### 3.5 Small-angle X-ray scattering

To further characterize the size of gp11 and obtain information regarding the shape of the quaternary gp11 assembly in solution, we conducted solution x-ray scattering experiments in the same Tris/NaCl buffer used in AUC experiments. No radiation damage was detected over the 10 s exposure time, so data obtained after the long exposure was used for analysis. The scattering data are not dependent on protein concentration, as the scattering curves superimposed well for protein samples at concentrations ranging from 1.4 to 2.7 mg/mL (figure 4).

**Figure 4:**
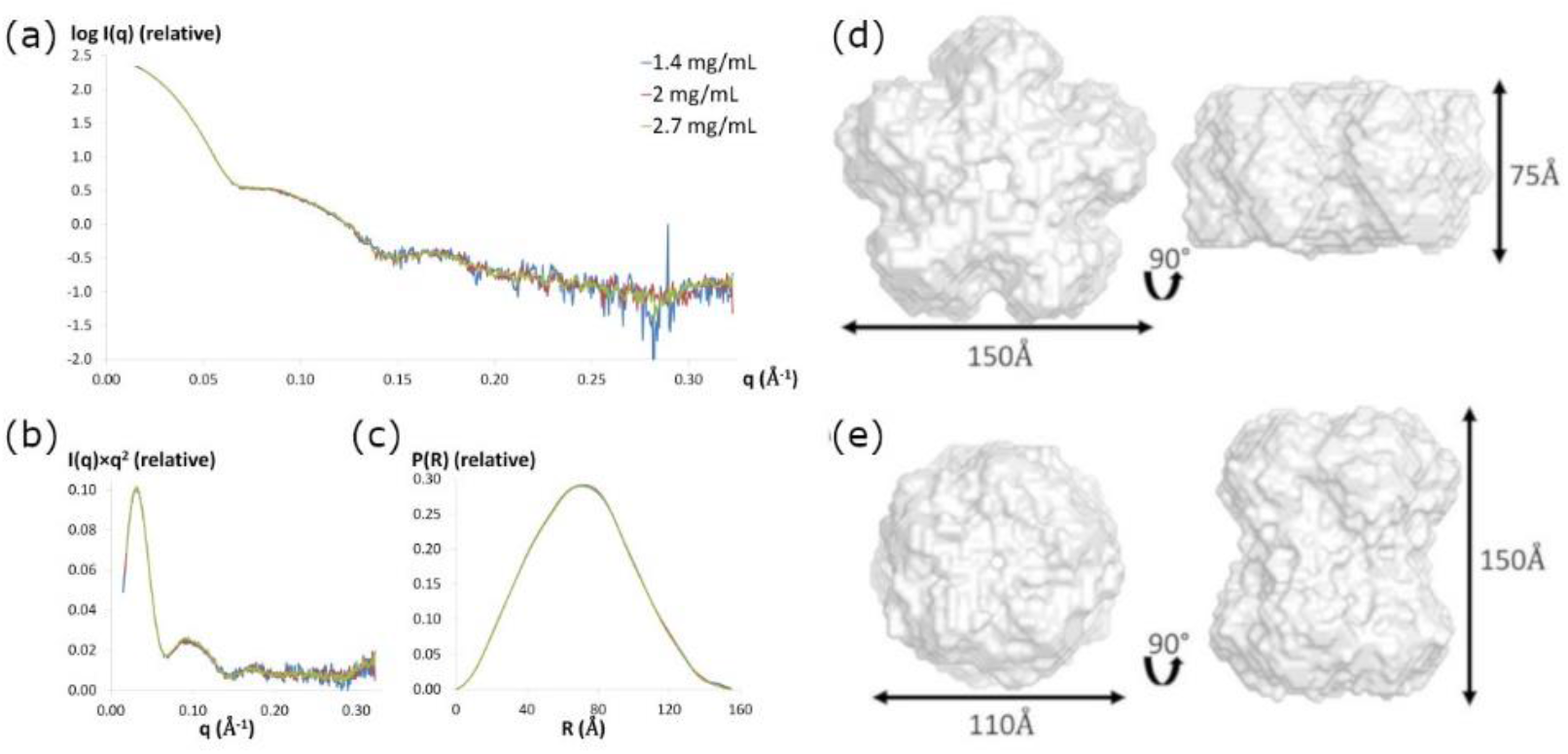
(a) Experimental scattering patterns for three concentrations of asccφ28 gp11 scaled to concentration. (b) Kratky plots and (c) pair-distance distance distribution functions P(R) of asccφ28 gp11 for three concentrations. Two classes of molecular envelopes, generated using GASBOR, are consistent with the X-ray scattering curve of gp11. D5 symmetry was imposed. Either a flatter oblate cylinder (d) or a more elongated prolate cylinder (e) are consistent with the SAXS measurements.

Two independent methods, the Guinier approximation and the pair-distribution function P(R), were used to calculate the radius of gyration (RG) value for each protein concentration. Both methods provided similar RG values, with ≈ 55 ± 2 Å calculated from the Guinier approximation and ≈ 54 ± 1 Å estimated from the pair-distance distribution function P(R) (Table 1). The maximum dimension of the particle, D_max_, was found to be 155 ± 5 Å by examining where the pair-distance distribution function went to zero. The hydrated particle volume (VP) was estimated between 745.6 and 778.6 nm^3^, corresponding to average molecular masses between 426.1 and 444.9 kDa, again indicating that gp11 forms a decameric assembly in solution (Table 1).

**Table 1:** SAXS parameters for the gp11 particle.

| Protein concentration | $R_G$ (Guinier) (Å) | $D_{\max}$ ($P(R)$) (Å) | $R_G$ (real space, $P(R)$) (Å) | Hydrated volume (Porod) (nm<sup>3</sup>) | Estimated molecular mass (kDa) | Number of Monomers |
| --- | --- | --- | --- | --- | --- | --- |
| 1.4 mg/mL | 54.8 | 155 | 53.8 | 745.6 | 426.1 | 9.5 |
| 2.0 mg/mL | 55.1 | 155 | 53.7 | 778.6 | 444.9 | 9.9 |
| 2.7 mg/mL | 55.2 | 155 | 53.6 | 760.6 | 434.6 | 9.7 |

### 3.6 SAXS ab initio shape calculations

The shape of the decamer in solution can be approximated from the shapes of the Kratky plot and the pair-distance distribution function, P(R). The Kratky plot (I(q)×q^2^ vs. q) (figure 4b) shows a bell-shaped peak at low angles that indicates a well-folded protein, and the pair-distance distribution function, P(R), shows a characteristic shape of a hollow globular, likely cylindrical, particle (Figure 4c) [39,47,48]. Eleven molecular envelopes of the decameric assembly were then obtained using the program GASBOR with D5 symmetry imposed on the SAXS data (X-ray diffraction results support this symmetry constraint, see results and discussion below). These molecular envelopes are indeed hollow cylinders, but segregate into two different classes with similar dimensions: an oblate shape for 6 models (model 1 ≈ 150 Å × 75 Å, figure 4d) and prolate shape for 5 models (model 2 ≈ 150 Å × 110 Å, figure 4e). Scattering curves from both classes of envelopes fit the experimental scattering data well, with χ2 values ranging from 1.1 to 1.2.

### 3.7 Electron microscopy

Negative stain images show a uniform collection of particles ~140-160 Å long and ~110-120 Å wide (figure 5). Most particles appear oblong, with an occasional nearly circular particle, presumably representing side and top views, respectively. All particles have a central dark spot, consistent with extra stain collecting in a void or channel. Based on the observed dimensions of side and top views and the accumulation of stain within particles, these results suggest a hollow cylindrical assembly, consistent with the prolate SAXS model.

**Figure 5:**
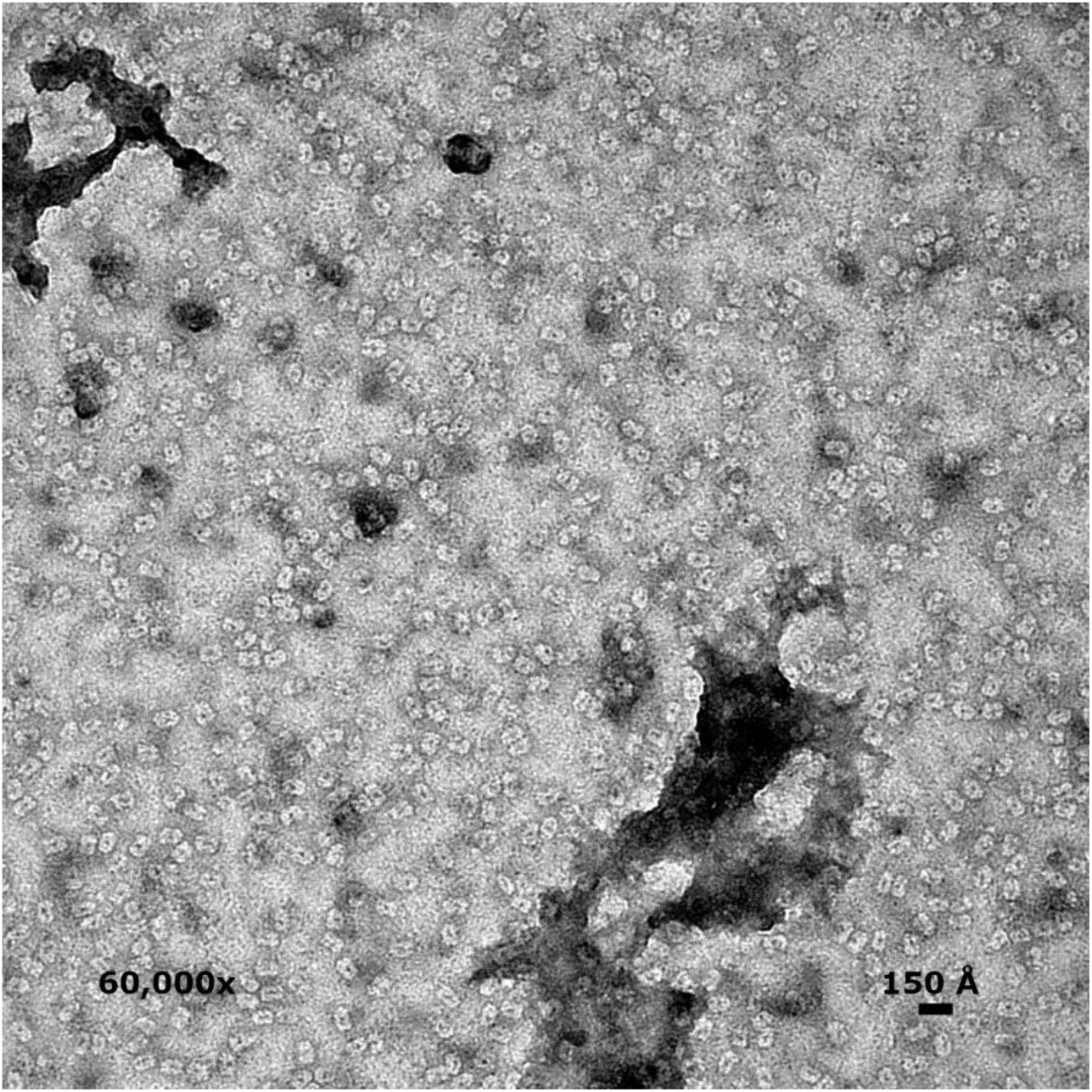
Uranyl acetate staining of gp11 showed cylindrical particles with heavily stained central regions. Magnification is 60,000x. These images more closely resemble the prolate envelope predicted by SAXS. A hollow cylinder would allow DNA to pass through the central channel of the motor.

### 3.8 Crystallization

Initial crystals of gp11 were obtained from the Wizard Classic screen (Rigaku) in 2 M ammonium sulfate, 0.1 M sodium citrate, pH 5.5 and Salt RX screen (Hampton Research) in 0.7 M citrate, 0.1 M Tris, pH 8.5. Gp11 has a predicted pI of 6.44, and thus would be positively or negatively charged, respectively, in the two conditions. Each condition served as the starting point for optimization, and crystals exceeding 100μm were eventually grown in 24-well VDX trays by hanging-drop vapor-diffusion over 1000 μL of well solution. A rhombohedral crystal (figure 6a) was grown over wells containing 1.5-1.8 M ammonium sulfate, 0.1 M citrate, pH 5.3-5.7. The drop consisted of 1μl protein at 4.3 mg/ml mixed with 1μl well solution containing 1.6 M ammonium sulfate, 0.1 M trisodium citrate, pH 5.7. The pH of trisodium citrate buffers was adjusted with hydrochloric acid. Crystals appeared after 2-4 days. Bi-pyrimidal crystals (figure 6b) were grown from basic conditions containing 1.0 M trisodium citrate, 0.1 M Tris pH 8.3. The final pH of this well solution was measured at 8.9. Protein concentration was 3.1 mg/ml, and 1.5 μL of protein was combined with an equal volume of well solution. Both solutions were pre-chilled, and the tray was set up and incubated at 277 K. Well-formed crystals typically took more than a month to grow.

**Figure 6:**
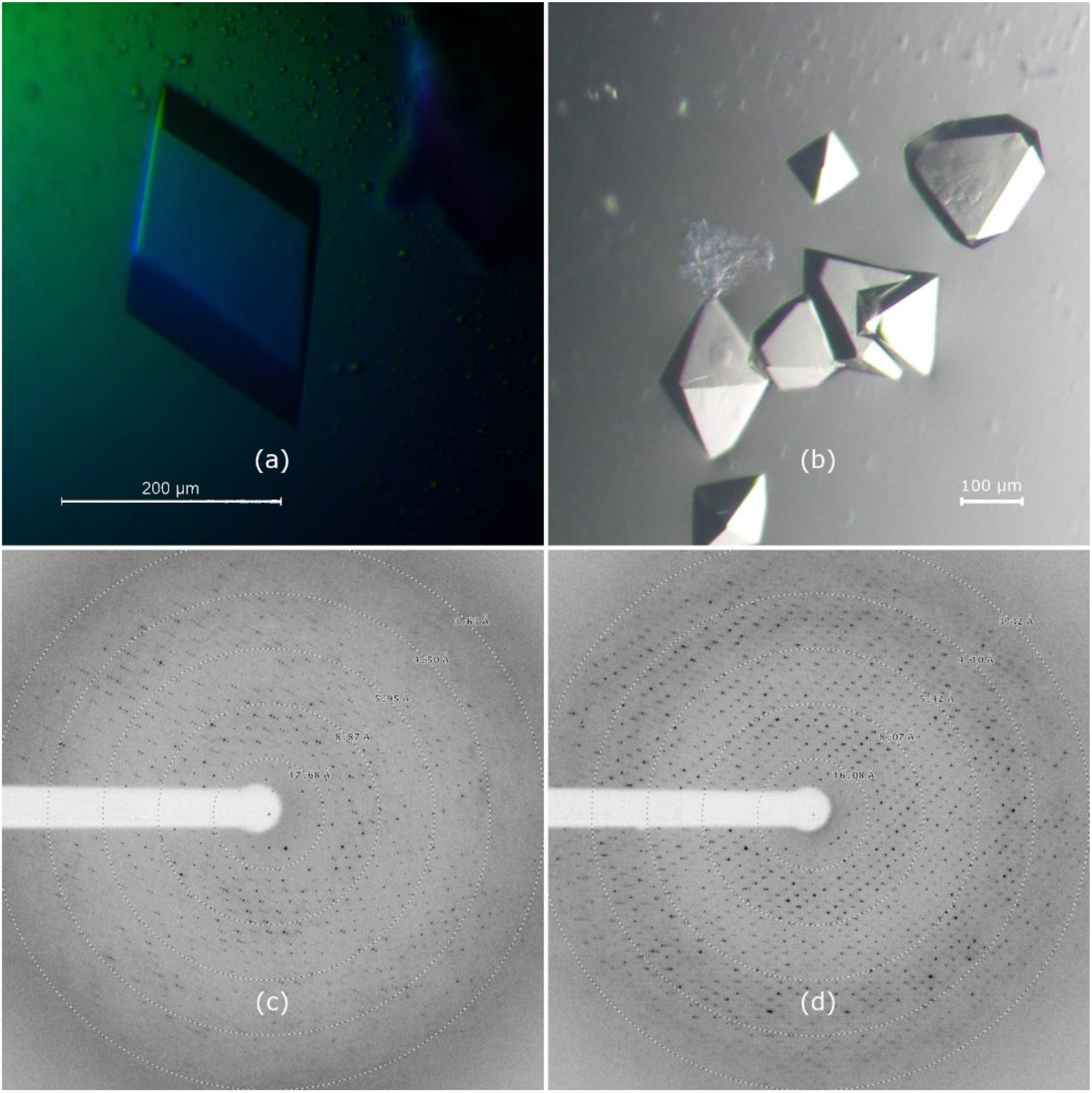
Crystals of gp11 grown under optimized conditions. (**a**) Crystal form A, grown in 1.6 M ammonium sulfate, 0.1 M trisodium citrate, pH 5.7. Crystals are rhombohedral with a maximum edge length of 200 μm. (**b**) Crystals form B, grown in 1.0 M trisodium citrate, 0.1 M tris(hydroxymethyl)aminomethane (Tris), pH 8.4. The truncated octahedron in the upper right measures approximately 150 μm x 150 μm x 50 μm. (c) Diffraction pattern for Crystal Form A, which indexed to space group P3_x_21. (d) Diffraction pattern for Crystal Form B, showing spots to ~2.8 Å, which indexed to P4_x_2_1_2.

### 3.9 Crystal X-ray diffraction

Diffraction data for optimized crystals were collected at Advanced Photon Source beamline 21 ID-F on a MAR 225 CCD detector. Diffraction patterns for each crystal are shown in figure 6, and data collection parameters summarized in table 2. Maximum diffraction of crystal form A extended to a resolution of ~3.3 Å. The crystal indexed as a primitive trigonal lattice type with unit cell dimensions a=b= 135.0 Å, c=76.7 Å. Systemic absences and merging statistics of symmetry related reflections were consistent with space group P3×21. The bi-pyramidal B-form crystals diffracted to 2.8 Å and were indexed as a primitive tetragonal lattice with unit cell dimensions a=b=110.9 Å, c=351.8 Å. Systematic absences and merging statistics of symmetry related reflections in this data set were consistent with space group P4_(1,3)_2_1_2. Matthews coefficient estimations of both crystal forms indicated a solvent content between 40 and 60%, corresponding to 4-7 ATPase copies in the asymmetric unit.

**Table 2:**
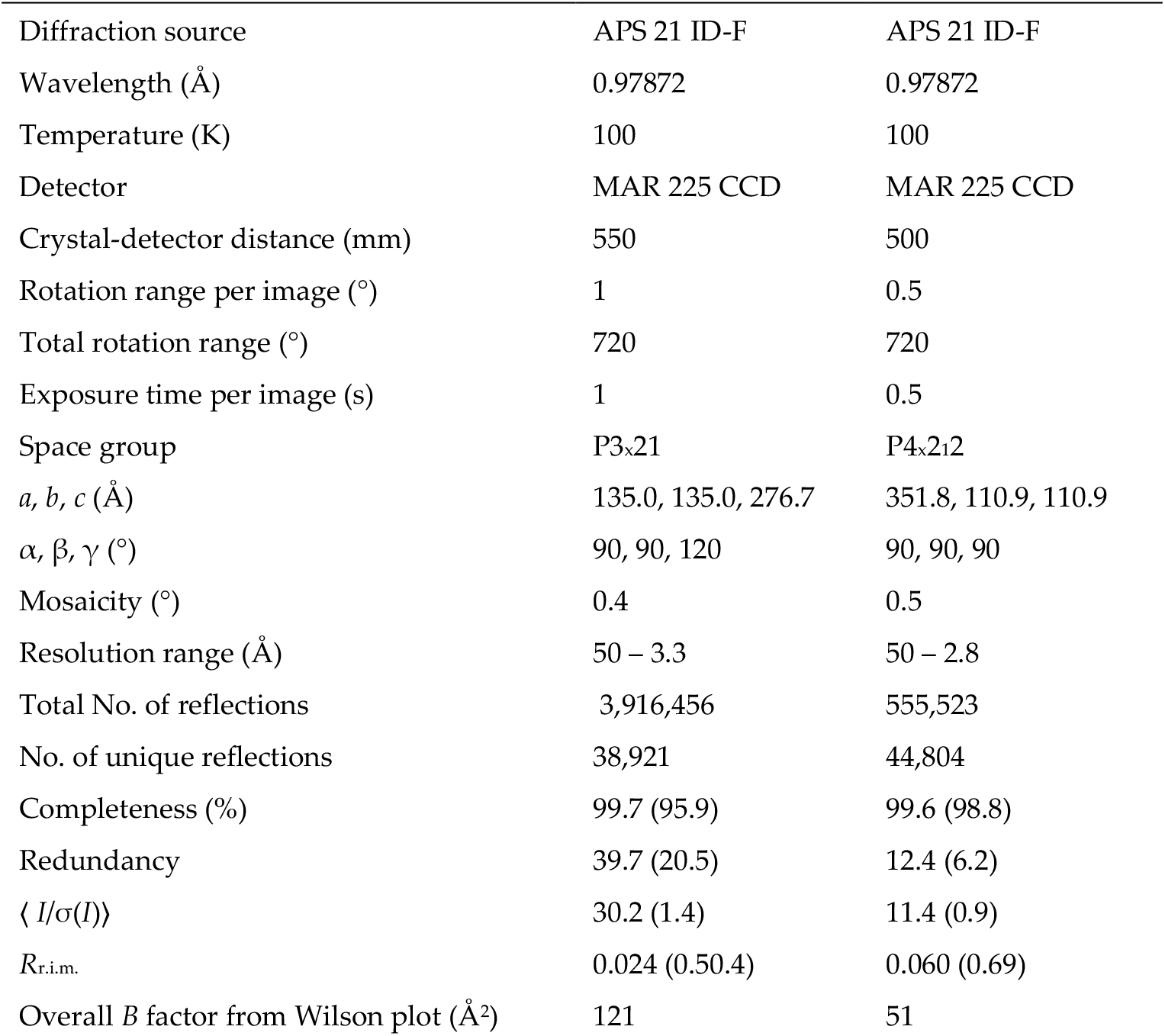
X-ray diffraction data collection and processing. Values for the outer shell are given in parentheses.

### 3.10 Activity assays

We measured ATPase activity for gp11 to assess whether the synthetic gene coded for an active enzyme product with the predicted NTPase activity. ATPase activity was determined by a coupled assay using the enzymes pyruvate kinase and lactate dehydrogenase, as shown in figure 7a. Reduction in absorbance at 340nm as a result of NADH depletion was monitored to assess the progression of the reaction. The change in absorbance units per unit time was obtained from the linear region of progress curves after steady state rates were observed for periods over five minutes. The values obtained from these slopes were then plotted as a function of substrate concentration (figure 7b) and the resulting curve was modeled by a nonlinear combination of the parameters V_max_ and K_m_ in the modified Michaelis-Menten equation:

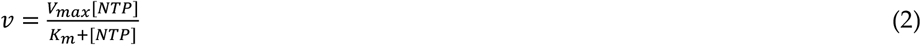

The values for V_max_ and K_m_ obtained from the nonlinear regression analysis were 7.6×10^-7^ Ms^-1^ and 28μM, respectively. Also, from V_max_=[gp11]k_cat_, k_cat_ was found to be 0.849 s^-1^.

**Figure 7:**
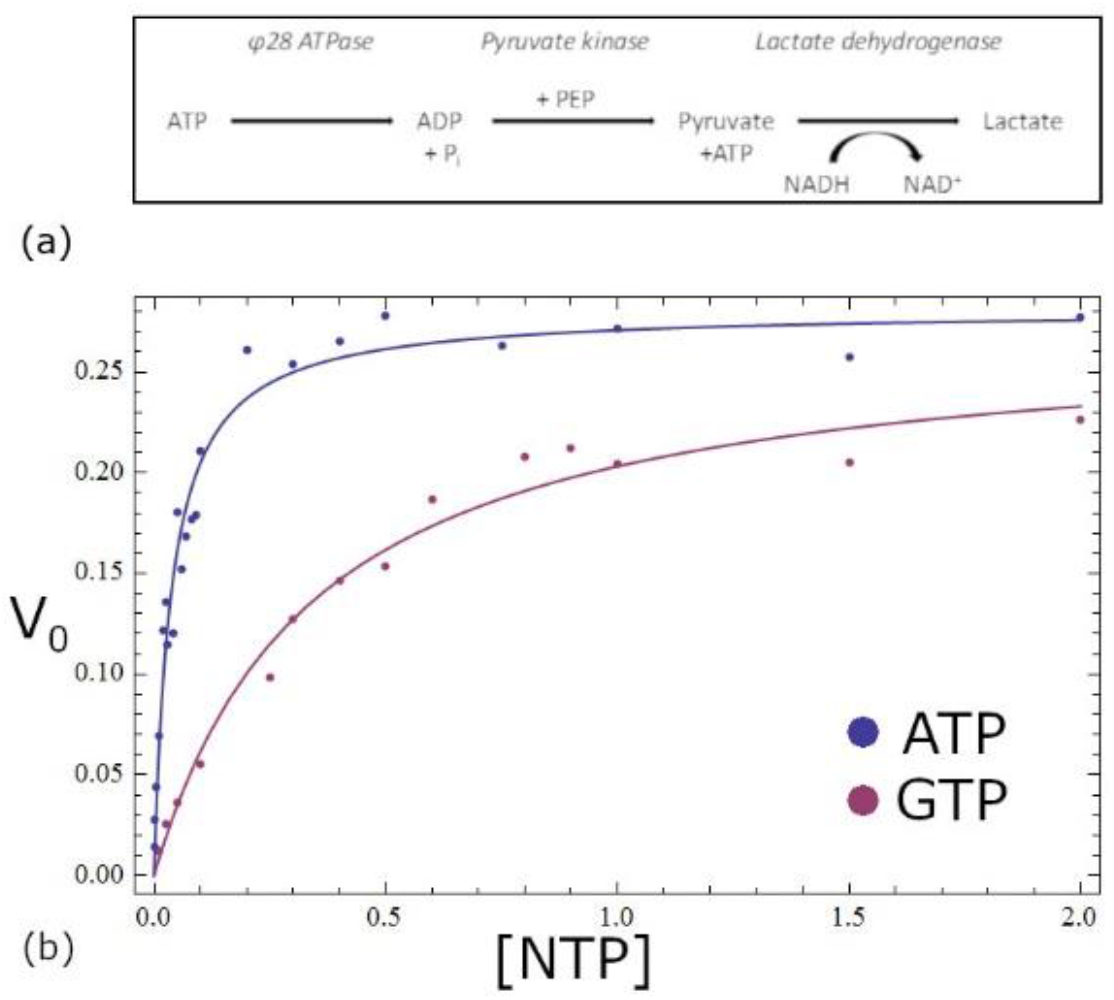
Steady-state kinetics for the asccφ28 encapsidation protein. (a) The gp11 ATPase produces ADP from ATP. Two subsequent enzymatic steps result in the conversion of NADP to NAD+, which is measured with a spectrophotometer. ATP hydrolysis by gp11 is the limiting reaction step, with all other reactants in large excess, so the ATP hydrolysis rate can be correlated with the measured reduction in NADP. (b) The upper blue plot represents ATP substrate, and lower magenta plot represents GTP. The 10-fold lower K_m_ for ATP indicates that gp11 is primarily an ATPase.

Since many NTPases are promiscuous regarding their selection of NTP substrates, we tested whether our construct was indeed primarily an ATPase. This is information is important since nucleoside triphosphate preference can offer clues as to the protein’s primary function. While there are many examples of GTPases within the NTP P-loop superfamily common to motor proteins, no GTPase has been reported to power genome encapsidation in phages, or in related motor proteins involved in nucleic acid recombination or unwinding [9]. The kinetic parameters found for GTP as a substrate were: K_m_=344 μM, V_max_=7.31×10^-7^ Ms^-1^ and kcat=0.817 s^-1^. Kinetic parameters for both substrates are summarized in Table 3. Based on the 10-fold increase in the Km value that occurred from switching from ATP to GTP, 28 μM to 344 μM, it seems that gp11 prefers ATP as a substrate.

**Table 3:** Kinetic parameters for the hydrolysis of NTP by gp11.

|  | ATP | GTP |
| --- | --- | --- |
| $K_m$ | $28 \mu\text{M}$ | $344 \mu\text{M}$ |
| $V_{\max}$ | $7.53 \times 10^{-7} \text{ Ms}^{-1}$ | $7.31 \times 10^{-7} \text{ Ms}^{-1}$ |
| $k_{\text{cat}}$ | $0.849 \text{ s}^{-1}$ | $0.817 \text{ s}^{-1}$ |

## 4. Discussion

Genome packaging motors have been studied extensively in several dsDNA bacteriophage systems. Proposed motor mechanisms invoke complex multi-step processes that require strict coordination between motor components to efficiently generate force. However, the molecular basis of subunit coordination and force generation remains unknown, largely due to challenges in structurally and biophysically characterizing the oligomeric ATPase ring motors that drive packaging. In currently available phage packaging systems, these ATPases typically have low solubility and are present only as monomers in solution. Thus, it is not possible to directly observe and interrogate the intermolecular interactions that regulate or coordinate ATP hydrolysis, force generation, or DNA translocation. Here, we report the biochemical, biophysical, and preliminary structural characterization of gp11, a DNA packaging ATPase from bacteriophage asccφ28.

Phage asccφ28 is genetically and morphologically similar to the well-studied bacteriophage φ29, yet there is no evidence for the unusual pRNA molecule that serves as a scaffold for assembly of a functional ATPase ring motor in φ29. Hence, we hypothesized that gp11 might assemble as functional ATPase rings in solution. Otherwise, asccφ28 is like φ29; gp11 and the asccφ28 portal protein are smaller and functionally simpler than their counterparts in genome-cutting packaging phages. Hence, asccφ28’s dsDNA encapsidation machinery may represent the minimum functional assembly of a viral dsDNA genome packaging motor. These properties, compositional simplicity and assembly of a functional ATPase ring motor, should facilitate detailed structural, biochemical, and biophysical analysis of a viral dsDNA packaging motor.

A recombinant, codon-optimized gene for gp11 was synthesized and expressed in *E. coli* with a C-terminal hexa-histidine tag to facilitate purification via metal ion affinity chromatography. After elution on a cobalt-containing Talon column, size-exclusion chromatography indicated that the protein assembled as a multimer in solution. Comparison to a standard elution curve of proteins with known molecular weight indicated a possible weight for the assembly ranging from 158 to 440 kDa. Subsequent sedimentation velocity and equilibrium analysis showed that gp11 is best represented as a single species in solution, with a molecular weight of ~445 kDa, which would correspond to a decameric assembly. Estimation of the Porod volume via small angle X-ray scattering confirmed this molecular weight, and further analysis of low angle scattering data provided additional shape information suggesting the assembly formed a cylinder, likely prolate in shape but possibly oblate. Negative stain EM allowed us to distinguish between these possibilities, clearly showing a prolate cylinder. Interestingly, while all biophysical mass and volume estimates pointed toward a decameric assembly, the dimensions of the asymmetric units in the two different X-ray crystallographic space groups could only accommodate between 4 and 7 copies of the ~45 kDa gp11 monomer. However, both space groups have crystallographic 2-fold axes of symmetry, suggesting that a molecular 2-fold symmetry axis is incorporated into global crystallographic symmetry. Hence, the simplest explanation parsimonious with the aggregate data is that gp11 is a decamer in solution, with approximate D5 molecular symmetry, i.e., an assembly consisting of two pentameric rings related by 2-fold symmetry.

While we cannot rule out the possibility that gp11 actually functions as a decamer when assembled on procapsids, we suspect that the dimerization of the two pentameric rings is an artifact of over–expression in the absence of asccφ28 procapsids. A significant argument against a biologically functional decamer is that the 2-fold symmetry axis relating the two pentameric rings requires that the two rings face opposite directions, either head-to-head or tail-to-tail. Hence, if both rings were to interact with DNA, the directions of imposed force would seem to cancel each other as the motor plays tug-of-war with the dsDNA substrate. Further, it is well established that the gp11 homologue in the closely related bacteriophage φ29 is a pentamer, as are the packaging ATPases in the more distantly related T4, T7, and P74-26 bacteriophages, suggesting that the biological assembly is a single pentameric ring.

Biochemical analysis of gp11 suggests that it binds and hydrolyzes ATP with affinities and rates similar to the functional ring forms of these other phage packaging ATPases. With the exception of bacteriophage lambda [27], recombinantly expressed packaging ATPases in other phages are monomeric, and are unable to bind and/or hydrolyze ATP when expressed recombinantly. Only upon oligomerization as pentameric rings on their respective procapsids are these enzymes able to bind and hydrolyze ATP. Hence the comparable ATP affinities and turn-over rates observed for recombinantly expressed gp11 suggests that the assembled pentameric rings reflect the biological assembly of the packaging ATPase on the procapsid.

## 5. Conclusions

Genome packaging motors have been studied extensively in several bacteriophage systems. Proposed motor mechanisms invoke multi-step processes that require strict coordination between motor components. However, the molecular basis of subunit coordination and force generation remains unknown, largely due to challenges in characterizing the ATPases that drive packaging. In currently available phage packaging systems, these ATPases typically have low solubility and are present as monomers in solution. Thus, it is not possible to directly observe and interrogate the intermolecular interactions that regulate or coordinate ATP hydrolysis and DNA translocation. Here, we report the expression and characterization of a DNA packaging ATPase from bacteriophage asccφ28 that self-assembles as a homomeric ring with high ATPase activity. These properties are promising for the detailed structural, biochemical, and biophysical analysis of a viral dsDNA packaging system.

## Author Contributions

Conceptualization, E.R.A., E.A.D., W.M.B., K.H.C and M.C.M; methodology, investigation and formal analysis, E.R.A., E.A.D., C.B., M.R.S., G.D., P.M., M.A.W and W.M.B.; writing E.R.A., E.A.D. and M.C.M; visualization, E.R.A and E.A.D.; supervision and funding acquisition, K.H.C and M.C.M.

## Acknowledgments

The authors acknowledge the Sealy Center for Structural Biology and Molecular Biophysics at the University of Texas Medical Branch at Galveston for providing research resources.

E.R.A. received a Houston Area Molecular Biophysics Training Program fellowship, funded by NIH grant T32GM008280 to Baylor College of Medicine, and a substantial portion of these experiments were complete as part of his Master’s degree work.

